# Hybrid Rank Aggregation (HRA): A novel rank aggregation method for ensemble-based feature selection

**DOI:** 10.1101/2022.07.21.501057

**Authors:** Rahi Jain, Wei Xu

**Affiliations:** Biostatistics Department, Princess Margaret Cancer Research Centre, Toronto, Ontario, Canada; Dalla Lana School of Public Health, University of Toronto, Toronto, Ontario, Canada

**Author notes:** Corresponding author (WX).

**Keywords:** high dimensional data, hybrid rank aggregation, artificial intelligence, machine learning, ensemble feature selection, random forest

## Abstract

**Background:** Feature selection (FS) reduces the dimensions of high dimensional data. Among many FS approaches, ensemble-based feature selection (EFS) is one of the commonly used approaches. The rank aggregation (RA) step influences the feature selection of EFS. Currently, the EFS approach relies on using a single RA algorithm to pool feature performance and select features. However, a single RA algorithm may not always give optimal performance across all datasets.

**Method and Results:** This study proposes a novel hybrid rank aggregation (HRA) method to perform the RA step in EFS which allows the selection of features based on their importance across different RA techniques. The approach allows creation of a RA matrix which contains feature performance or importance in each RA technique followed by an unsupervised learning-based selection of features based on their performance/importance in RA matrix. The algorithm is tested under different simulation scenarios for continuous outcomes and several real data studies for continuous, binary and time to event outcomes and compared with existing RA methods. The study found that HRA provided a better or at par robust performance as compared to existing RA methods in terms of feature selection and predictive performance of the model.

**Conclusion:** HRA is an improvement to current single RA based EFS approaches with better and robust performance. The consistent performance in continuous, categorical and time to event outcomes suggest the wide applicability of this method. While the current study limits the testing of HRA on cross-sectional data with input features of a continuous distribution, it could be applied to longitudinal and categorical data.

## Introduction

Feature selection (FS) is important for high dimensional data analysis [1,2] as it helps in reducing the feature space which mitigates the issues related to model fitting, generalizability [3] and computational complexity [4,5]. Many techniques exist in literature to conduct FS, but the commonly used methods can be broadly classified into base feature selection and ensemble-based feature selection (EFS). The FS techniques in the base feature selection are sub-classified into Filter, Wrapper, and Embedded approaches [6,7]. The internal structure of features in a dataset is evaluated for filter-based FS [8–10]. Multiple subsets of features are evaluated to select the best subset of features for wrapper-based FS. In the case of the embedded-based FS approach, the final model itself performs feature selection by penalizing the excess features during the model building process [11–13].

In the EFS approach, features are selected based on the performance across multiple models rather than one model prepared using the base feature selection approach. EFS normally performs better than the base approach [14]. EFS is a multi-step approach where literature provides different techniques for each step (Figure 1). The first step creates multiple datasets from the original data either by sampling of features [15], samples [16,17] or both [18,19], or by duplicating the original dataset [15,20,21]. Each of these datasets can be processed incorporating interaction terms for the features [19] and undergoing feature transformation [18]. The second step develops a model for each of the datasets created in the first step by using either a single base technique or multiple base techniques. The approach which uses datasets sampled from the original dataset and a single base technique to build the model is called the homogenous EFS approach [22]. The approach which uses the original dataset and multiple base techniques to build the model is called the heterogenous EFS approach [20]. The third step is the extraction of feature performance from each of the models using either single [16] or multiple metrics [17]. The feature performance is pooled or aggregated in the next step and is referred to as rank aggregation (RA). Finally, the features are selected based on their pooled performance or importance cut-off [15].

**Figure 1:**
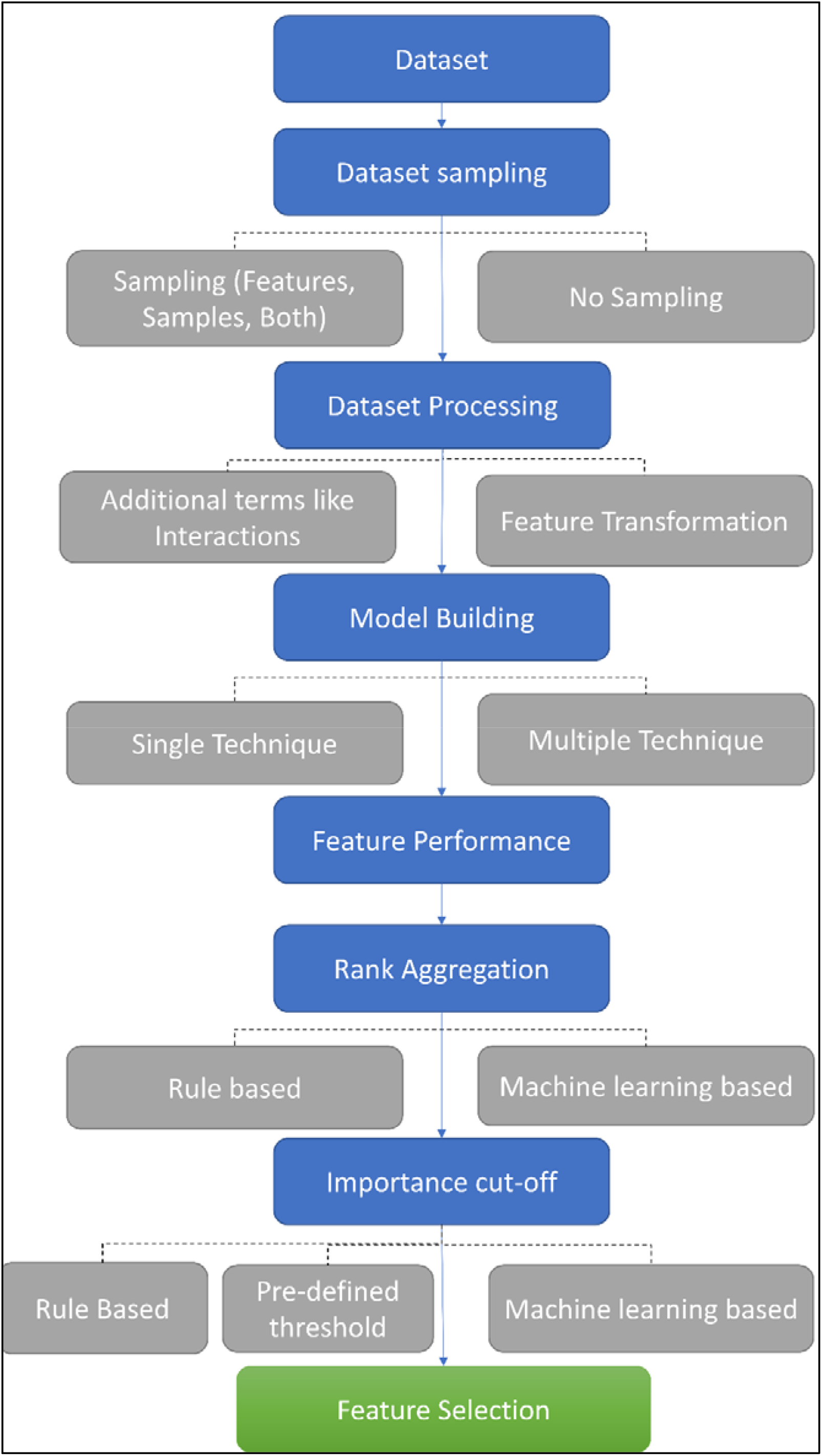
Ensemble based feature selection approach.

RA technique has a strong influence on the EFS approach performance. The RA techniques used in the literature could be classified into two categories namely rule-based RA and machine learning-based RA. The rule-based RA techniques pool the feature performances based on a pre-defined rule which prevents them from dynamically learning and creating rules based on the data. Some of the commonly used techniques to aggregate the feature performance are frequency [23,24], mean, median and robust rank aggregation (RRA) [25,26]. In the machine learning-based RA techniques, the rules are created dynamically by the technique based on the dataset [27]. Supervised RIDGE is one such ML-based RA technique which provides pooled feature performance by estimating its influence on the study objective using RIDGE regression [27]. However, no single RA technique could give consistently best performance in all scenarios. Thus, multiple RA techniques need to be evaluated to both determine the best RA technique for a study and determine expected features and their performance in a study.

It is desirable to develop a RA approach which is more robust than the existing RA techniques. Accordingly, this study proposes a novel RA method called hybrid rank aggregation (HRA) which generates multiple rankings or pooled performances of a feature from different RA techniques. Then, unsupervised learning-based feature selection is performed to select features based on performance across multiple RA techniques. The proposed HRA-based EFS is novel in many ways. Firstly, it uses a unique perspective which creates a shift from a single RA technique per EFS algorithm to multiple RA techniques per EFS algorithm. Secondly, it provides an approach to select features based on their performance across multiple RA techniques rather than a single RA technique. Thirdly, it allows the usage of both rule-based RA and ML-based RA techniques in a single EFS algorithm. Fourthly, the usage of unsupervised learning-based technique at the feature selection step allows the algorithm to have rules which can dynamically adapt to the dataset. Finally, its ability to integrate with existing ensemble methods makes it a versatile approach which enhances the applicability of existing EFS methods.

This paper is divided into the following sections. The “Methodology” section explains the HRA-based EFS whose performance is evaluated and compared with existing RA methods using simulations in the “Simulation Studies” section. Further, in the “Real Studies” section, HRA is tested in several real data to determine its practical relevance. Finally, the paper is summarized with future research directions in the “Conclusion and Discussion” section.

## Methodology

HRA methodology is developed to pool results from different RA techniques in the rank aggregation step of the ensemble learning for robust results (Figure 2). EFS process is initiated for a dataset containing *n* samples, *p* features, and an outcome. Initially, multiple models are created by either a homogenous or heterogeneous approach. Then, RA is performed using different RA techniques to obtain multiple feature rankings. An unsupervised learning algorithm selects features that are important across multiple RA techniques. The proposed methodology is discussed in detail below.

**Figure 2:**
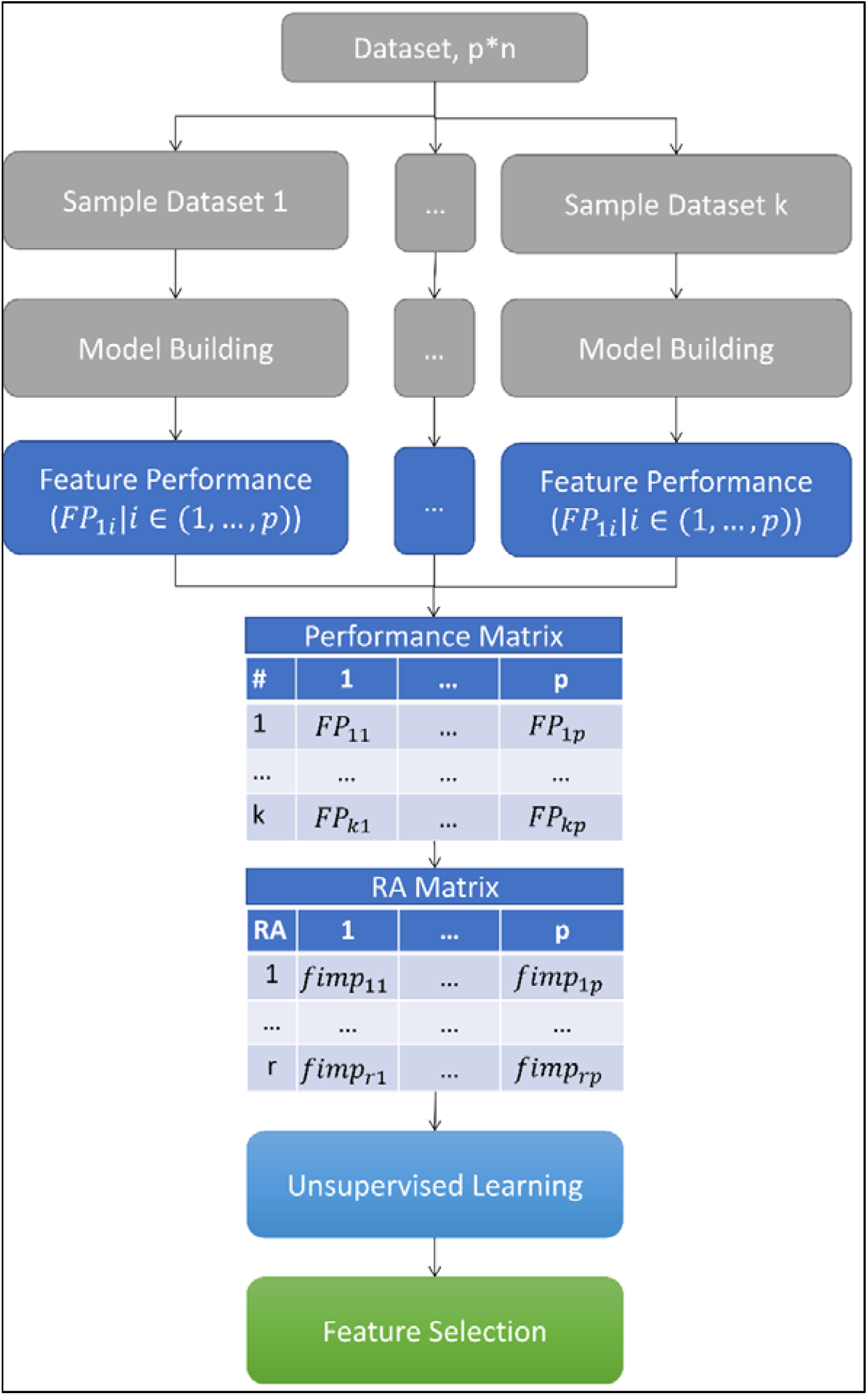
Graphical representation of HRA-based ensemble feature selection

### Generate multiple models

A model is prepared for each of the *k* sample datasets created from the original dataset *D* containing *p* features and *n* samples for an outcome *y* by randomly sampling *j*_*i*_ | *i* ∈ {*1*, …, *k*},*1* < *l* ≤ *p* features from *p*. Each of the sample datasets contains *n* samples obtained by subsetting samples from *D* with replacement. For *k* sample datasets, *m* models are created using any modeling technique.

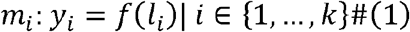

where function *f* for *i*^*th*^ dataset depends upon the technique used to build the model *m*_*i*_. In the current study, the random forest technique is used for building *m* models.

### Feature performance matrix

The feature performance *FP* of all features for *m* models is obtained to create a feature performance matrix *FP*_*mat*_.

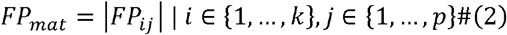

The different metrics are used as *FP* in literature like a coefficient estimate of feature [15,16], goodness of fit [18], feature importance [28] and feature significance [21]. For illustration purposes, feature importance *a* is used as *FP*. Since, in any model, only *t*| *t* ≤ *p* features are used, thus many cells in the *FP*_*mat*_ will not have a *a* value. In such cases, cells without *a* value are either dropped from the analysis or replaced by values zero based on the RA technique used in the next step.

### Rank Aggregation

Literature suggests different RA techniques to aggregate the *FP*_*mat*_ to obtain the feature importance *fimp* of *p* features from *m* models [25,26]. *fimp* is generated for *r* RA techniques to create *RA*_*mat*_.

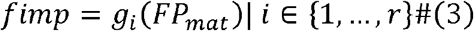

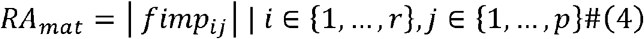

Where, *g*_*i*_ is the RA function used to get *fimp* of *p* features for *i*^*th*^ RA technique. Different RA techniques used to generate *fimp* are mean based RA (MeRA), maximum based RA (MaRA), minimum based RA (MiRA), median based RA (MedRA), coefficient of variation based RA (CVRA), standard deviation based RA (SDRA), robust rank aggregation (RRA), t-test based RA (tRA), Wilcoxon signed-rank test based RA (WRA), supervised LASSO (SL), supervised RIDGE (SR) and supervised random forest (SRF) [27]. The supervised learning-based RA techniques namely SL, SR and SRF also need a label, hence predictive performance of *m* models on the samples left out of the bootstrap samples are used as the label values [27].

### Feature Selection

In the conventional approach where single RA technique is used to generate *fimp* of *p* features from *m* models, machine-learning [19], rule-based methods [29] and pre-defined threshold-based methods [18,19] exist which defines a *fimp* cut-off to select the final features *q*_*best*_. In the HRA approach, features are selected based on their *fimp* across multiple RA techniques. Similar to the conventional approach, HRA assumes that the more important features are relevant for building the model. Thus, features with higher *fimp* across the different RA techniques are chosen as selected features. To dichotomize *p* features into high and low *fimp* groups, unsupervised clustering is performed. Weighted K-means clustering is used in this study to cluster *p* features based on their *fimp* values across *r* RA techniques given in *RA*_*mat*_. The features present in the cluster with higher mean *fimp* value is selected as final features *q*_*best*_.

In the case of weighted K-means clustering, all RA techniques are not given equal importance to avoid excessive influence of weak performing RA techniques on the outcome. Thus, *r* RA techniques are given weights *w =* {*w*_*l*_,…, *w*_*r*_} based on their performance. However, RA techniques performance is unknown apriori, so *w* weights have to be either provided arbitrarily [17] or need to be derived from existing information. For illustration purpose, the current study derives *w* weights from existing information. In each of the *r* RA techniques, *p* features *fimp* values are obtained followed by splitting the *p* features based on their *fimp* values into two groups using K-means clustering. The features in the group with high mean *fimp* values are selected and called as selected features *q*_*r*_ for the *r*^*th*^ RA technique. These *q*_*r*_ features are used to train a predictive model on the dataset *D* and predictive performance is obtained and is used as weight *w*_*r*_for the *r*^*th*^ RA technique. For illustration purpose, RIDGE technique is used to build predictive model and 10-fold cross-validation with three repeats is performed to get the predictive performance. The current study allocated *w*_*r*_ weights exponentially to *fimp* values of *R*_A_ techniques to get weighted *fimp* values for *r* RA techniques 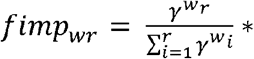, where *y* is the user-defined exponential factor. Pseudo Algorithm given below summarizes the complete HRA-based EFS algorithm.

#### Pseudo Algorithm: HRA-based ensemble feature selection

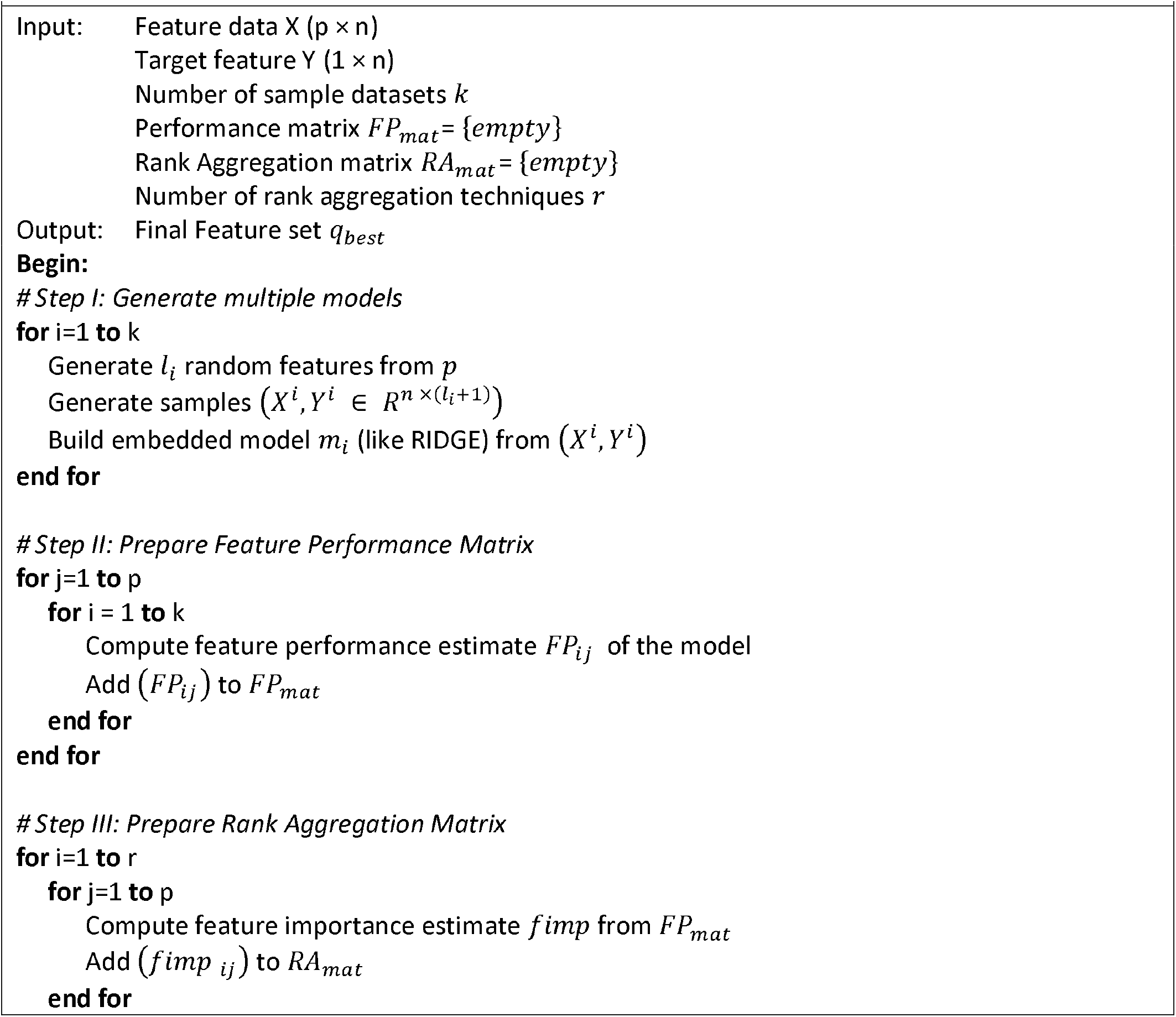
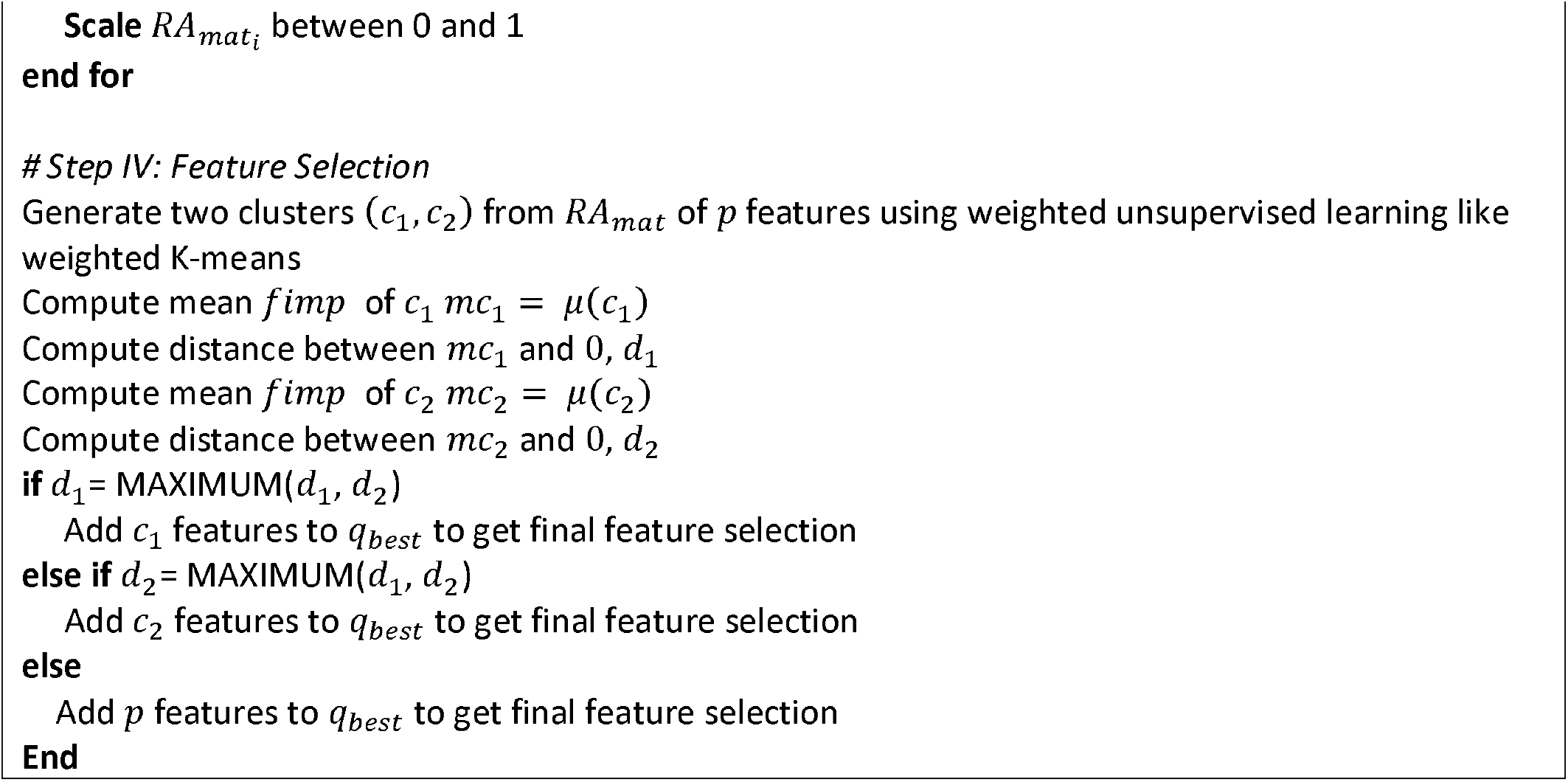

### Simulation Studies

The study generates multiple simulated scenarios to evaluate the proposed HRA method and compare its performance with other RA methods using a regression underline model 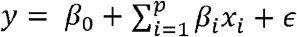 for a continuous outcome with effect size *β*, features *x* ∼*N*(*0, 1*) and error term *ε*∼*N*(*0, σ*^*2*^)| *σ = 0*.*25*. 500 random scenarios are generated by changing *p, n, µ*, the number of features with multicollinearity and the covariance value between multicollinear features (Table 1). Non-zero *β* is assigned to the true features only.

**Table 1:** Description of the simulation data

| # | Parameter | Range |
| --- | --- | --- |
| 1 | Number of features, $p$ | {50,75,100} |
| 2 | Number of samples in training data, $n$ | {50,75} |
| 3 | Non-zero coefficient, $\beta$ | {-0.50, 0.50, 0.75} |
| 4 | Number of true features | [5,20] |
| 5 | Number of features with non-zero correlation with each other | {15,20,25} |
| 6 | Covariance value of features with non-zero correlation with each other | {{-5,5}, {-8, 8}} |
| 7 | Number of Bootstraps | 50 |

A random forest-based homogenous ensemble approach is used for the study. 50 models are created by creating subsets of data containing 2 to *p* features and n samples obtained after resampling from the original dataset with replacement. In the HRA method, final feature selection is obtained by pooling feature ranks obtained from 12 different RA methods namely, mean based RA (MeRA), maximum based RA (MaRA), minimum based RA (MiRA), median based RA (MedRA), coefficient of variation based RA (CVRA), standard deviation based RA (SDRA), robust rank aggregation (RRA), t-test based RA (tRA), Wilcoxon signed-rank test based RA (WRA), supervised LASSO (SL), supervised RIDGE (SR) and supervised random forest (SRF). During the pooling stage, the feature ranking is normalized. Among the RA methods giving identical feature ranking, one RA method is randomly selected while other RA methods are dropped from the pooling step. Further, RA methods giving the identical rank to all features are also dropped from the pooling step. The remaining RA methods are pooled using weighted K-mean clustering.

The performance of the proposed HRA method is compared with the abovementioned 12 existing RA methods. Inbuilt packages of statistical language R 4.0.3 are used for performing analysis. Penalized regression like LASSO and RIDGE is performed using *glmnet* package [30], random forest model is prepared using *randomForest* package [31] for continuous and binary outcome and *randomForestSRC* for survival outcome and RRA is performed using *RobustRankAggreg* package [26].

The performance of different RA methods is compared using the following three metrics. The first metric is the ability of different models to discriminate between true features and noise features which is measured using the F1 Score. The second metric is the predictive performance of the features selected by a RA method. The RIDGE based predictive model is built using the training data. The study generated a test dataset of 500 samples to get the predictive performance measured using inverse root mean square error (RMSE). The third metric measures the percentage of target features retained in the final list of selected features.

500 simulated scenarios are generated to compare HRA and existing RA methods. It is found that the performance of RA methods varied with the scenario, which suggests that internal data characteristics could play a role in determining the appropriate RA method (Table 2). In the case of F1 metric, it is found that MeRA and HRA gave the best discrimination ability compared to other RA methods. However, MeRA is not good at selecting the target features, rather it was MaRA and HRA that are good in selecting the target features. In the case of the predictive performance of selected features, it is found that HRA gave the best performance. Overall, the results suggest that HRA is a robust RA approach for feature selection as it consistently gave better or at par performance than existing RA methods. Thus, HRA could be a good candidate to select the target features. Further, HRA could help in improving the performance of existing ensemble-based approaches for high-dimensional data by performing better RA as compared to existing methods.

**Table 2:** Comparison of HRA method with Existing RA methods in terms of target feature selection, feature discrimination ability (F1 Score) and outcome prediction (1/RMSE) for 500 scenarios

| RA technique |  | Scenarios |  |  |
| --- | --- | --- | --- | --- |
|  |  | Target Features<br>(% of total target features) | F1 Score | Predictive Performance (1/RMSE) |
| | | $\mu$[95%CI] | | |
| **Existing** | CVRA | 66.85 [64.58-69.13] | 0.46 [0.45-0.48] | 1.11 [1.06-1.16] |
|  | MARA | **88.94 [87.83-90.05]** | 0.59 [0.57-0.6] | 1.62 [1.56-1.68] |
|  | MeRA | 71.28 [69.46-73.09] | **0.69 [0.68-0.71]** | 1.36 [1.3-1.42] |
|  | MedRA | 60.21 [58.28-62.15] | **0.66 [0.64-0.67]** | 1.18 [1.12-1.23] |
|  | MIRA | 41.78 [39.8-43.77] | 0.52 [0.50-0.54] | 0.92 [0.87-0.97] |
|  | RRA | 56.02 [53.94-58.11] | 0.61 [0.59-0.62] | 1.1 [1.05-1.15] |
|  | SDRA | 2.58 [2.00-3.15] | 0.02 [0.02-0.03] | 0.34 [0.31-0.36] |
|  | tRA | 66.03 [63.74-68.32] | 0.46 [0.45-0.48] | 1.11 [1.06-1.17] |
|  | WRA | 43.8 [42.49-45.12] | 0.25 [0.24-0.26] | 0.66 [0.64-0.69] |
|  | SL | 20.84 [19.31-22.37] | 0.29 [0.27-0.3] | 0.63 [0.6-0.66] |
|  | SRF | 29.01 [27.69-30.34] | 0.26 [0.25-0.27] | 0.66 [0.64-0.69] |
|  | SR | 45.09 [43.4-46.78] | 0.53 [0.51-0.54] | 0.92 [0.88-0.97] |
| **HRA** | RF | **85.15 [83.58-86.73]** | **0.65 [0.64-0.67]** | **1.68 [1.61-1.74]** |

### Real Studies

The performance of HRA and existing RA methods are also compared in four real studies. Study I uses US county data (p = 578 and n=3141) related to non-communicable diseases which is obtained from Community Health Status Indicators (CHSI) study [32]. Study II uses US elderly data (p=1470 and n=4377) related to their health and wellbeing which is obtained from National Social Life, Health and Aging Project (NSHAP) study [33]. Study III and Study IV are genomic datasets. Study III contains data on DNA methylation (p = 27578 and n = 108) with human age as the outcome [34,35]. Study IV contains gene expression data (p = 56602 and n = 476) of the prostate cancer patients obtained from TCGA [36]. The outcome of the first three studies is continuous, while in Study IV a binary and a time-to-event outcome are considered. IDC status is used as the binary outcome and Progression-Free Survival (PFS) is used as the time-to-event outcome of Study IV.

The dataset for all the studies is cleaned and processed for ease of analysis (Table 3). During this processing, features and samples are screened to remove highly correlated features, non-continuous features, missing values and features with very low standard deviation. The final cleaned datasets are split randomly into training and testing datasets. Feature selection and predictive model is prepared using the training data while test data is used to measure the predictive performance. Inverse RMSE, area under curve (AUC) of receiver operating characteristics (ROC) and Harrell’s C-index are used as the predictive performance metrics for models depending upon their outcome. The RA methods are compared based on their mean performance across ten trials. For each study, 100 ensemble models are prepared and 10 trials are performed.

**Table 3:** Summary of the real datasets

| Real Studies | Marginal Features (p) | Outcome feature(s) | Sample size (n) |  |  | k |
| --- | --- | --- | --- | --- | --- | --- |
|  |  |  | Total | Train | Test |  |
| *Study I* | 80 | Health Status | 97 | 77 | 20 | 100 |
| *Study II* | 45 | Height | 1035 | 207 | 828 | 100 |
| *Study III* | 2289 | Age | 108 | 86 | 22 | 100 |
| *Study IVa* | 500 | IDC Status | 476 | 380 | 96 | 100 |
| *Study IVb* | 500 | Progression Free Survival (PFS) | 472 | 377 | 95 | 100 |

The results suggest that the HRA method provided a robust performance that is better or at par with existing RA methods (Table 4). The stable HRA performance suggests that it might be a reliable approach to get target features as compared to single metric RA methods. Further, it is found that the performance of HRA prevailed in the models with binary and survival outcomes, which suggests that HRA could perform in different data types. Thus, HRA seems to be a more robust approach than existing RA methods. While, in certain scenarios, some existing methods could give better performance than HRA but overall, the existing methods are less robust than HRA in giving good results.

**Table 4:**
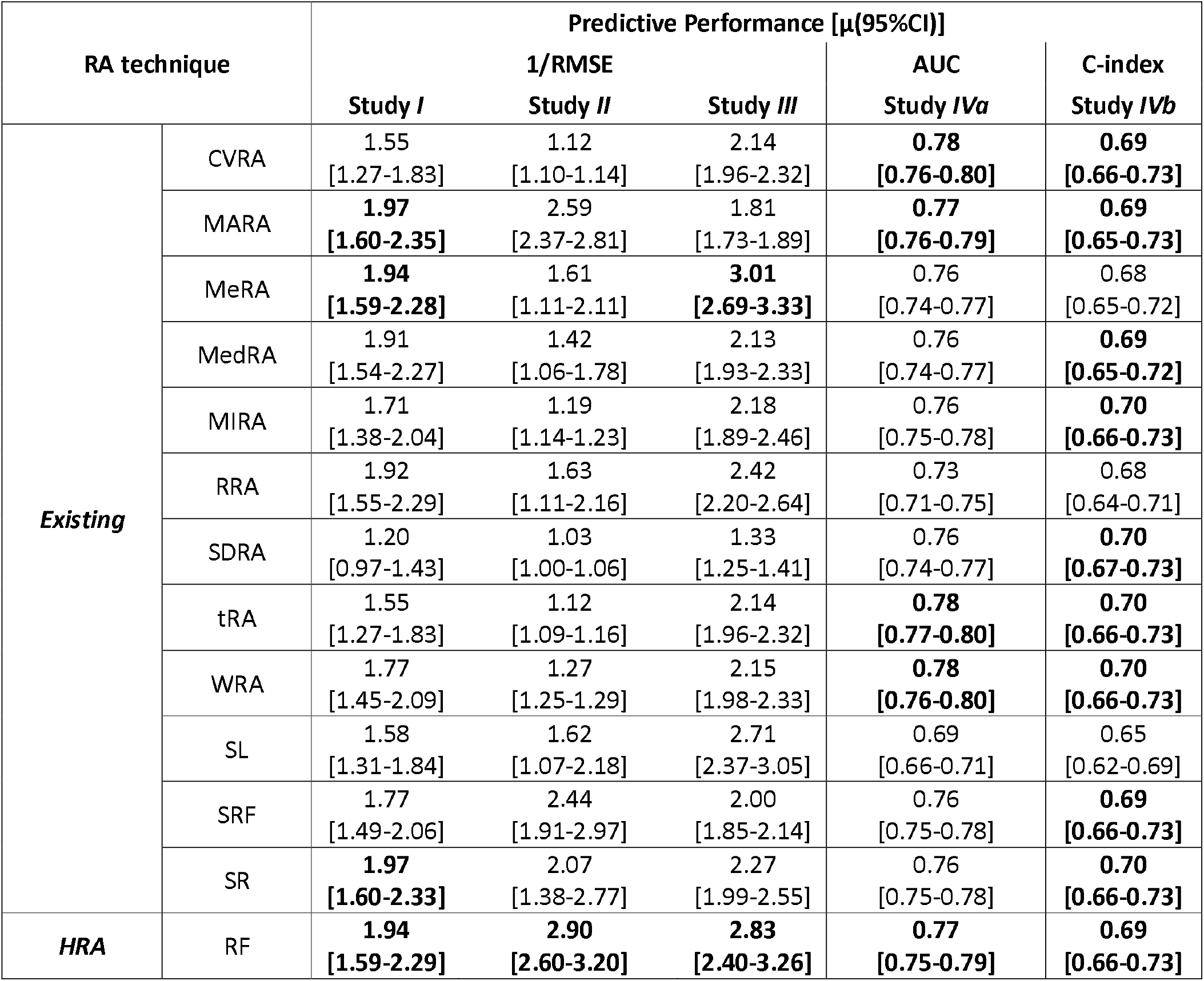
Comparison of SRA methods with Existing methods for three real study scenarios in terms of outcome prediction (1/RMSE)

## Conclusion and Discussion

This paper proposes an innovative rank aggregation method, HRA, for the EFS. The HRA approach pools the feature importance from the existing RA methods to select features using weighted unsupervised clustering. This approach is flexible as it could work with a wide variety of ensemble techniques, data types and RA methods. Moreover, HRA can select true features with good discrimination between target and noise features. The simulated studies and real dataset studies show the consistent and good predictive performance of the approach. The good performance in real studies suggests the practical relevance of the approach.

HRA still has certain limitations which need to be addressed in future studies. Firstly, the current scope is limited to concept testing. Thus, future studies could evaluate different techniques to build ensemble models and perform unsupervised clustering to determine the influence of technique on HRA performance. Secondly, HRA is tested with models having a linear combination of true features, it will be relevant to determine the HRA performance in models with a non-linear combination of true features.

## Declarations

### Ethics approval and consent to participate

Not Applicable

### Consent for publication

Not Applicable

### Availability of data and materials

All the datasets and code are in the github link: https://github.com/rahijaingithub/HRA.

## Author Contributions

ALL AUTHORS HAVE READ AND APPROVED THE MANUSCRIPT.

**Conceptualisation:** RJ, WX

**Formal Analysis:** RJ

**Investigation:** RJ

**Methodology**: RJ, WX

**Software:** RJ

**Supervision:** WX

**Validation:** RJ, WX

**Writing-original draft:** RJ

**Writing-review & editing:** RJ, WX

## Acknowledgements

Not Applicable

